# A combination of two-enzyme system and enzyme engineering improved the activity of a new PET hydrolase identified from soil bacterial genome

**DOI:** 10.1101/2024.02.01.578500

**Authors:** Hideaki Mabashi-Asazuma, Makoto Hirai, Shigeru Sakurai, Keigo Ide, Masato Kogawa, Ai Matsushita, Masahito Hosokawa, Soichiro Tsuda

## Abstract

We here report a novel PET hydrolase originating from a soil microbial genome sequence. This enzyme, bbPET0069, exhibits characteristics resembling a cutinase-like Type I PET-degrading enzyme but lacks disulfide bonds. Notably, bbPET0069 displayed remarkable synergy with *Candida antarctica* lipase B (CALB), demonstrating rapid and efficient PET degradation. To improve the PET degradation activity of bbPET0069, we employed a three-dimensional (3D) structural modeling to identify mutation sites around its substrate binding domain combined with a protein language model for effective mutation prediction. Through three initial rounds of directed evolution, we achieved a significant enhancement in PET degradation with CALB, resulting in a 12.6-fold increase compared to wild-type bbPET0069 without CALB. We confirmed its PET degradation activity in PET nanoparticles and films, and our proposed approach enabled efficient PET degradation to terephthalic acid monomers up to 95.5%. Our approach, which integrates a two-enzyme system with protein engineering, demonstrates the potential for enhancing the activity of emerging PET-degradation enzymes, which may possess unique attributes.

**Graphical Abstract:** A novel PET hydrolase, bbPET0069, was identified from a soil microbial genome. bbPET0069 and CALB showed remarkable synergy in PET degradation. Using surface feature analysis, PET degradation activity of bbPET0069 was significantly improved. This combination of a two-enzyme system and surface feature analysis holds promise for enhancing emerging PET-degradation enzymes.

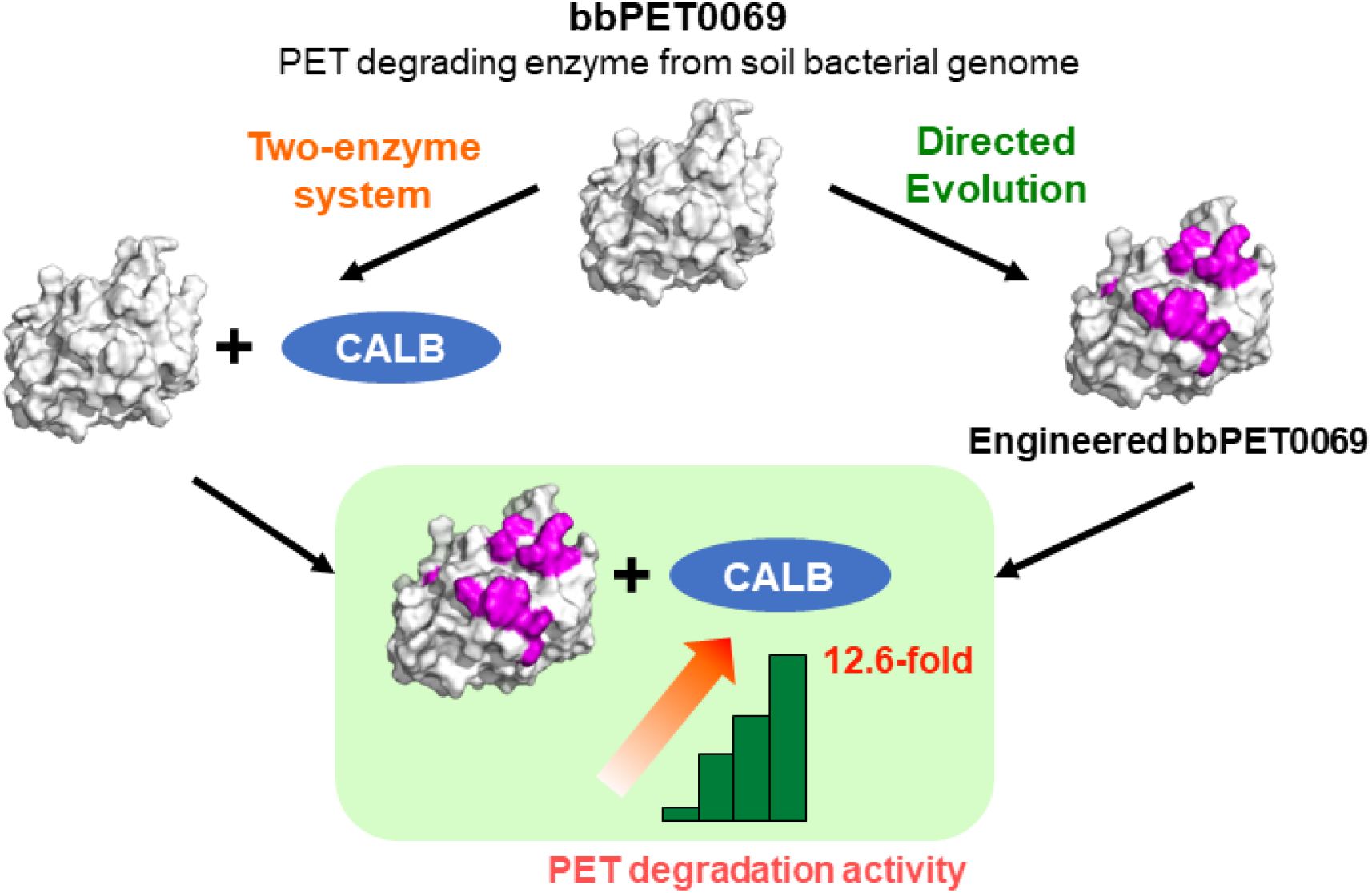

## Introduction

In recent years, the issue of plastic waste has gained widespread recognition as a global concern. According to data from the United Nations, approximately 460 million tonnes of plastic are produced annually worldwide, with only 9% being recycled ^[1]^. Most of this plastic waste is either disposed of in landfills, incinerated, or persists in environmental settings such as oceans, contributing to detrimental impacts on the ecosystem. A substantial amount of plastic waste has been recycled using material recycling methods and several recycling approaches, including chemical recycling ^[2]^. However, a shared limitation among these methods is the necessity for high-temperature treatments surpassing several hundred degrees Celsius. Thus, there is an increasing need to advance environmentally friendly recycling techniques.

Bio-based plastic degradation through enzymatic processes and the subsequent utilization of its decomposition byproducts represent a promising recycling method, offering an alternative to chemical recycling that circumvents the necessity for high-temperature treatments. This approach involves converting polymers into monomers, holding considerable promise as a method requiring lower temperatures for plastic breakdown ^[3,4]^. The most advanced research in enzyme-mediated plastic degradation is polyethylene terephthalate (PET). Following the initial discovery of TfH as a PET-degrading enzyme ^[5]^, subsequent findings of potent PET-degrading enzymes such as LCC ^[6]^ and IsPETase ^[7]^ have led to the development of numerous variants showing enhanced PET degradation activity and improved thermal stability ^[8–13]^. Although preprocessing of PET is required, this enzymatic approach to plastic recycling is poised to be the pioneering example of practical application ^[14]^.

Improving the efficiency of enzymatic PET degradation is essential to make enzymatic plastics recycling a viable option. Current efforts in protein engineering of PET degrading enzymes have often focused on improving thermal resistance to create enzymes capable of withstanding temperatures around the glass transition temperature (Tg) of PET, which typically ranges between 60-70 °C ^[15]^. Conversely, it is also suggested that PET degradation is feasible regardless of Tg, as long as the enzyme exhibits robust activity ^[16]^. Among various strategies to enhance PET degradation, the synergistic use of PET-degrading enzymes in combination with other enzymes has also increased the amount of PET degradation. One approach involves fusing IsPETase and IsMHETase via a linker peptide, resulting in efficient PET degradation ^[17]^. The other way was mixing the PET-degrading enzyme HiC with CALB, possessing BHET and MHET degradation activity ^[18]^. Recent reports suggest an enhanced activity of PET-degrading enzymes when combined with engineered enzymes possessing BHET degradation activity ^[19]^. Thus, exploring strategies that augment degradative potential by combining multiple enzymes, rather than solely focusing on enhancing the activity of PET-degrading enzymes, proves to be an effective approach.

In this study, we identified a new PET degrading enzyme, bbPET0069, and assessed boosting PET degradation using a two-enzyme system and protein engineering hybrid strategy. Notably, bbPET0069 displayed favorable synergistic behavior with CALB. The coexistence of both enzymes significantly increased PET degradation in a CALB concentration-dependent manner. We applied directed evolution and a novel surface feature analysis tool based on estimated protein structures to enhance its activity. This tool pinpointed amino acid positions likely to increase activity. Introducing mutations at these sites notably improved PET degradation. Ultimately, the mutant enzymes retained beneficial synergy with CALB, resulting in a considerable increase in PET degradation. This combined approach, integrating CALB and surface feature analysis-based mutagenesis, represents a novel strategy for rapidly enhancing PET degradation, particularly under ambient reaction conditions.

## Results and Discussion

### Identification of bbPET0069 as a Functional PET Hydrolase

We first searched our in-house microbial genomic database, called bit-GEM, using a well-characterized PET hydrolase, Leaf-branch Compost Cutinase (LCC), as a reference. This led to the identification of a functional PET degrading enzyme among twelve sequences similar to LCC. This soil microbe-derived sequence, bbPET0069, displayed complete conservation of the catalytic triad (S132, D178, H210), which are contained in all PET-degrading enzymes. Among known PET-degrading enzymes, bbPET0069 exhibited the highest amino acid identity with PHL7 classified under Type I, reaching 56.0% identity. Furthermore, it displayed identities of 44.1% and 41.0% with LCC and IsPETase, classified as Type I and Type IIb enzymes, respectively (Fig. 1A). Furthermore, in bbPET0069, the residue x1 in the G-x1-S-x2-G motif, commonly observed in all PET-degrading enzymes, was histidine instead of tryptophan, aligning with the characteristic features of Type I enzymes. Based on these findings, bbPET0069 is closely related to the Type I of the PET-degrading enzyme family based on its primary sequence. Interestingly, due to the absence of cysteine residues at the presumed positions for a disulfide bond in bbPET0069, it is assumed that this enzyme likely lacks any intramolecular disulfide bonds. Typically, Type I PET hydrolases possess one pair, while Type II PET hydrolases possess two disulfide bonds ^[20]^. Next, the protein structure of bbPET0069 was estimated using AlphaFold2 ^[21]^. The result revealed an α/β fold, a typical protein structure in all PET-degrading enzymes. The estimated structure of bbPET0069 was aligned well with the structure of PHL7 ^[22]^ (RMSD = 0.539) (Fig. 1B).

**Figure 1.**
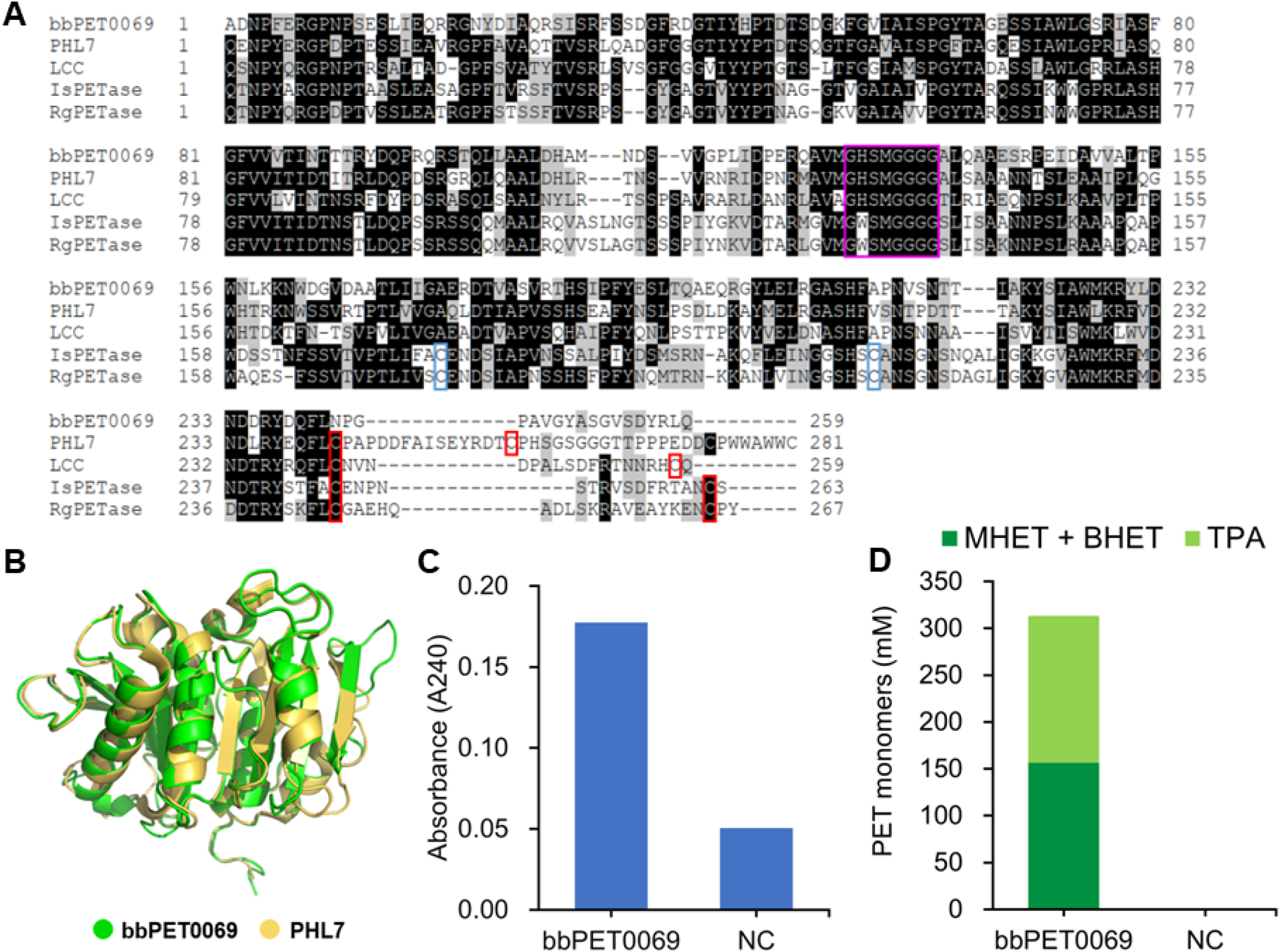
Identification of bbPET0069 as a functional PET hydrolase. A) Sequence alignment of bbPET0069 and the previously known PET hydrolases, including PHL7, LCC, IsPETase, and RgPETase. The purple box shows the G-x1-S-x2-G motif found in all PET hydrolases. Red boxes are the disulfide bonds shown in Type I and Type II. Blue boxes are the disulfide bond shown in only Type II enzymes. B) Structural alignment of the bbPET0069 structure predicted by AlphaFold2 and PHL7. C) PET hydrolase activity of bbPET0069 determined by the increase of A240. NC shows no enzyme control. D) PET degradation products (MHET, BHET, and TPA) produced by bbPET0069. NC was no enzyme control, and the level of the peaks was under the detection limit of our standard curve (labeled as the asterisk).

First, we examined ester bond cleaving activity in bbPET0069 by incubation assay with para-nitrophenyl butyrate (pNPB). Although it showed weaker activity than the LCC F243I mutant, which showed strong PET-degrading activity ^[9]^, we confirmed esterase activity in bbPET0069 (Supporting Fig. 1).

Next, to measure PET degradation activity, the purified enzyme was reacted with amorphous PET film (crystallinity of 8.7%). As a result, a clear increase in absorbance at 240 nm, indicative of PET monomers, was observed compared to the negative control without the enzyme (Fig. 1C). To confirm that the increase in absorbance at 240 nm indeed originated from PET degradation products such as terephthalic acid (TPA), mono-(2-hydroxyethyl) terephthalic acid (MHET), and bis-(2-hydroxyethyl) terephthalic acid (BHET), the reaction products of the PET film were analyzed by HPLC. The result showed that the reaction solution of bbPET0069 and PET film contained approximately equal amounts of TPA and MHET plus BHET (Fig. 1D). Thus, bbPET0069 was confirmed to be a functional PET-degrading enzyme. bbPET0069 exhibited PET degradation activity at 40 °C. However, at temperatures of 50 °C and above, it lost its activity (Supporting Fig. 2). One possible reason for the lack of thermal resistance in bbPET0069 might be the absence of any intramolecular disulfide bonds within its structure.

**Figure 2.**
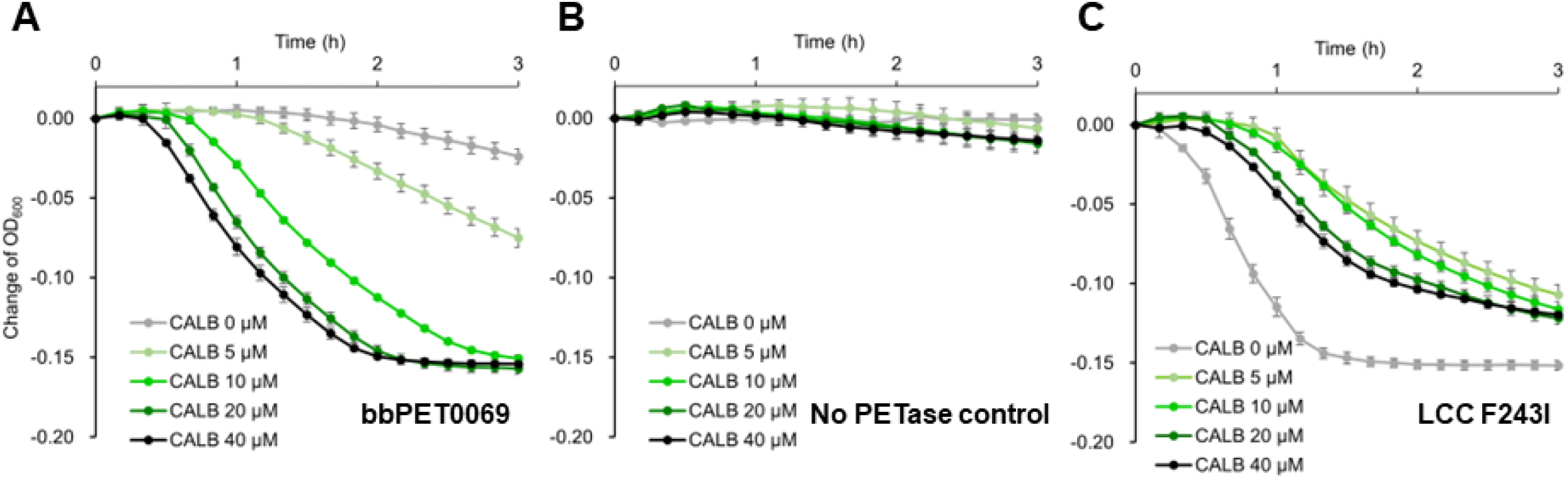
PET nanoparticle degradation assays. A) PET nanoparticle degradation assay performed with 500 nM of bbPET0069 with or without CALB. B) PET nanoparticle degradation assay performed with different concentrations of CALB. C) PET nanoparticle degradation assay performed with LCC F243I mutant with or without 20 µM CALB.

### Two-Enzyme System with CALB Boosted the PET Hydrolase Activity of bbPET0069

To investigate whether mixing bbPET0069 with another enzyme enhances PET degradation activity, PET nanoparticle degradation assays were conducted by adding various concentrations of CALB to bbPET0069 because this assay is much faster and sensitive than PET film degradation assay ^[23]^. In all experiments, purified bbPET0069 was used at 0.5 µM. The results revealed a concentration-dependent escalation in PET degradation rates with CALB. The most rapid PET degradation was observed upon the addition of 40 µM CALB, and the highest level of PET degradation was confirmed at 20 µM and 40 µM CALB (Fig. 2A). The most significant increase in PET degradation activity induced by CALB was observed after 110 minutes of the reaction, showing a 90-fold increase in PET degradation in the presence of 40 µM CALB compared to its absence.

To exclude the possibility that CALB itself degraded most of the PET, PET nanoparticle degradation assays using various concentrations of CALB in the absence of bbPET0069 were performed. The results showed a slight decrease in turbidity when adding ten µM or more of CALB compared to the absence of CALB (Fig. 2B). However, this decrease was much smaller than the degradation activity observed with 0.5 µM bbPET0069 alone (Fig. 2A, Gray line). Therefore, these results indicated a substantial increase in PET degradation activity due to the combination of bbPET0069 and CALB.

Although there have been multiple reports of combining CALB with PET-degrading enzymes at similar concentrations ^[17,18,24,25]^, attempts to add CALB at 10-40 times higher concentrations than the PET-degrading enzyme have not been made. Moreover, the discovery that excess CALB remarkably increased the PET degradation quantity of the PET-degrading enzyme by about 6.4 times in 3 h is a novel observation from this experiment.

Subsequently, we investigated whether the increase in PET degradation quantity due to CALB could apply to known PET-degrading enzymes. CALB was added to one of the highly active LCC mutants, LCC F243I. Contrary to bbPET0069, the presence of CALB was found to be inhibitory against the PET degradation activity of LCC F243I (Fig. 2C). Particularly, a significant reduction in PET degradation quantity was observed in the presence of 10 µM CALB. However, no correlation was observed between the added CALB concentration and PET degradation activity. Similarly, when adding CALB to FAST-PETase, another potent PET-degrading enzyme, the PET degradation quantity was significantly suppressed (Supporting Fig. 3). It is postulated that LCC rapidly degrades MHET to TPA during the PET degradation process, potentially reaching the endpoint, and even with the addition of CALB, no further changes might occur. Additionally, there could be mutual compatibility between the PET-degrading enzymes where the effects of CALB addition are seen. As each PET-degrading enzyme has different optimal reaction buffers and pH levels, the buffer used when CALB was added might have influenced the activity of each PET-degrading enzyme.

**Figure 3.**
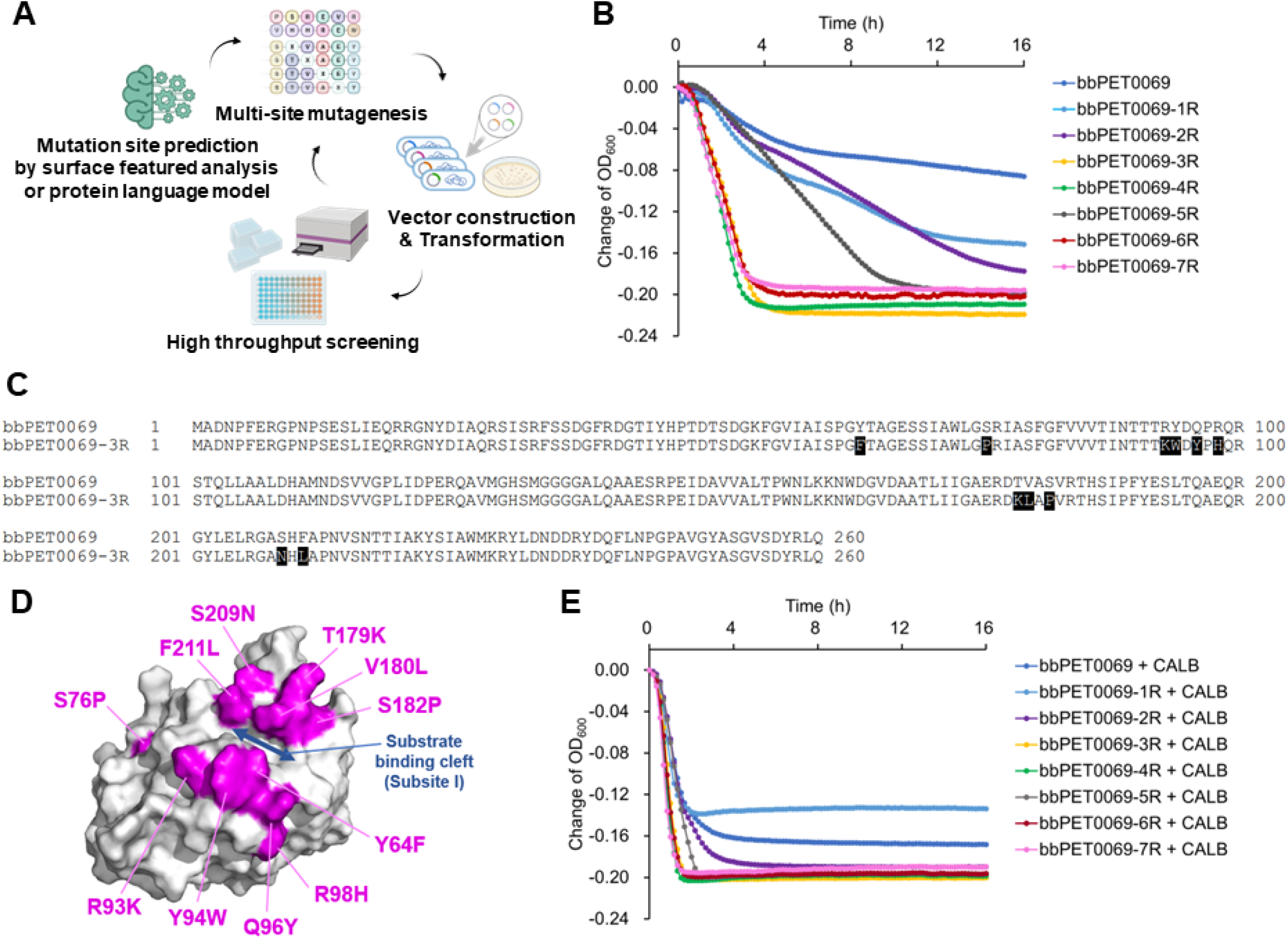
bbPET0069 mutants produced through our directed evolution campaign. A) Our directed evolution workflow. B) PET nanoparticle degradation assay using the best mutant from each round of mutagenesis. C) Amino acid sequence alignment between wild-type bbPET0069 and bbPET0069-3R. The eleven mutation sites were colored as black. D) The AlphaFold2 structure model of bbPET0069-3R. The mutated amino acids were labeled as purple. The PET substrate binding cleft, subsite I, was labeled as a blue arrow. E) PET nanoparticle degradation assay using bbPET0069 and its mutants with 20 µM CALB.

### Directed Evolution of bbPET0069 to Enhance its PET Hydrolase Activity

Next, we set out to conduct a total of seven rounds of enzyme engineering by directed evolution. We employed three approaches to introduce mutations: (1) a 3D structural modeling, called Surface Feature Analysis where we computationally identified amino acids exposed on and around the surface of active site, (2) Computational prediction using protein language models for stabilizing mutations, and (3) incorporating mutations previously identified as beneficial mutations in known PET-modified enzymes. These methods used for introducing mutations in each round are presented in Supporting Table 1. Each round involved identifying target mutation sites and amino acid residues following different rules, followed by cloning via PCR to create a library of mutant enzyme-expressing *Escherichia coli*. The PET degradation activity of these *E. coli* lysates was measured using a PET nanoparticle turbidity reduction assay ^[23]^ for all candidates (Fig. 3A).

As a result, a significant increase in activity was observed in the best mutant of the third round, namely bbPET0069-3R (Fig. 3B).

For the first and second rounds, mutation sites were predicted using the Surface Feature Analysis around the substrate binding site. Based on these predictions, multiple mutations were introduced simultaneously, creating mutant libraries that were then screened. Both bbPET0069-1R (S76D) and bbPET0069-2R (S76P/R93K/Y94W/Q96T/R98F) showed superior PET degradation compared to the wild-type. However, no significant change in the initial degradation rate was observed (Fig. 3B - Blue, light blue, and purple). In the third round, a total of 11 mutations (Y64F/S76P/R93K/Y94W/Q96Y/R98H/T179K/V180L/S182P/S209N/F211L) that had exhibited increased activity in the previous rounds were incorporated into bbPET0069-2R. As a result, bbPET0069-3R showed a substantial increase in activity compared to bbPET0069-2R (Fig. 3B - Yellow). bbPET0069-3R exhibited a 5-fold increase in activity compared to the wild-type after 110 minutes of reaction and proceeded to completely degrade PET nanoparticles in the solution in about 4 hours. In the fourth round, we used a protein language mode, ESM-1v, to predict the mutation sites that would likely improve the stability of the enzyme^[26]^. The screening results showed that the top mutant of the fourth round, bbPET0069-4R, was faster than bbPET0069-3R (Fig. 3B - Green). In the fifth round, multiple mutants were created by combining mutated sites showing improved degradation activity in the fourth round and compared for their PET degradation. However, even the most active variant in this round, bbPET0069-5R, showed a significant decrease in activity compared to the best clone of the fourth round, bbPET0069-4R (Fig. 3B -Black). It suggests that while the mutations tested in the fourth round were calculated based on the sequence of bbPET0069-3R, combining them does not necessarily lead to increased activity. Therefore, in the sixth round, the sequence of bbPET0069-5R was not used, and instead, bbPET0069-4R was employed as the template sequence, utilizing the new version of the protein language model ESM2 ^[27]^. Similar to the fourth round, scores were computed for each amino acid residue when substituted with other amino acids, determining which amino acid to mutate based on these scores and constructing a mutant library. However, from this library, no mutants demonstrating activity exceeding bbPET0069-4R were obtained (Fig. 3B - Red). In the final seventh round, several mutants were created by introducing single mutations into positions corresponding to the amino acids that have been tested in previously reported PET-degrading enzyme mutants, known to contribute to increased activity and stability ^[8–13,28–32]^. As a result, only the R184S mutation showed a slightly faster degradation than the sixth mutant but was very similar to bbPET0069-4R (Fig. 3B - Pink). The mutations in positions corresponding to this amino acid were found in N212K of HotPETase and R228S in the Cut190 mutant.

In summary, significantly increased activity was observed in the third-round mutants, contributing significantly to the enhanced activity throughout the seven rounds (effectively six rounds, considering the failure of the fifth round). The mutant library from the third round was generated by combining mutations determined to improve activity in the first and second rounds through surface feature analysis. Thus, it might be crucial to efficiently combine the proposed mutation sites and mutated amino acids, focusing on the substrate-binding region highlighted in surface feature analysis, compared to the protein language models used in the fourth and sixth rounds, to obtain highly active mutant variants. Investigating whether this mutation introduction method is similarly effective in other enzymes would be intriguing, representing a compelling avenue for future studies.

### Directed Evolution of bbPET0069: Comparing the Mutated Amino Acids with the Previously Reported PET degrading enzymes and their mutants

When comparing the mutation sites and the number of mutations between the most active mutants in each mutation round and the wild-type sequence of bbPET0069, it was observed that the mutant exhibiting the highest activity enhancement, bbPET0069-3R from the third round, had mutations in 11 amino acid positions (see Supporting Table 2). On the other hand, despite introducing mutations in 20 amino acids in bbPET0069-7R, a significant increase in activity was not achieved. It suggests that for bbPET0069, optimizing activity while maintaining protein stability might be achieved with around 11 amino acid mutations. Moreover, bbPET0069-3R was developed by exploring 1068 mutants in the first, 1068 in the second, and 712 combinations in the third. Considering the entire process of directed evolution in this study, it can be concluded that at least two rounds of mutation collection and one round of combination were suitable for enhancing the activity of bbPET0069.

When aligning the amino acid sequences of the wild-type bbPET0069 and bbPET0069-3R, it appears that the 11 amino acid mutations are dispersed throughout the entire sequence (Fig. 3C). However when the amino acid sequence of bbPET0069-3R was predicted using AlphaFold2, it became evident that, except for S76P, the remaining ten mutations are situated on both sides flanking the substrate-binding cleft (Fig. 3D). Comparing the estimated structure of bbPET0069 to the reported structure of PHL7, which exhibited the highest sequence similarity to bbPET0069 ^[22]^, it was observed that among these ten positions, four (Y64F, Y94W, Q96Y, V180L) were within subsite I, and six (R93K, R98H, T179K, S182P, S209N, F211L) were located within two amino acids of subsite I (Supporting Table 3). Conversely, subsite II of bbPET0069 demonstrated conservation similar to PHL7 and LCC, particularly in the extended loop region where the amino acids were well conserved only with LCC. Some identical amino acid mutations that matched those found in LCC (F64Y, Q96Y) or PHL7 (F211L) were observed. L210, especially in PHL7, emerged as a highly crucial amino acid for its strong PET degradation activity, a conclusion supported by our independent analysis through a different approach, which we find remarkably intriguing ^[33]^.

In the case of Type I enzymes such as LCC and PHL7, mutations have frequently been designed rationally ^[31,34,35]^. Conversely, several reports used directed evolution for Type IIb enzymes like IsPETase. These include DepoPETase, which combines seven mutations ^[13]^; DuraPETase, a combination of 10 mutations ^[10]^; and HotPETase, integrating 21 mutations after seven rounds of mutagenesis ^[11]^. When mapping these mutation sites onto the crystal structure of IsPETase, unlike the mutations introduced in bbPET0069-3R by us, the positions of mutations are observed throughout the entire protein, not just within the substrate-binding region (Supporting Fig. 4). It remains challenging to provide a clear explanation for the substantial increase in PET degradation activity resulting from mutations in amino acids that do not belong to a limited number of ligand-binding sites, as seen in enzymes like FAST-PETase.

**Figure 4.**
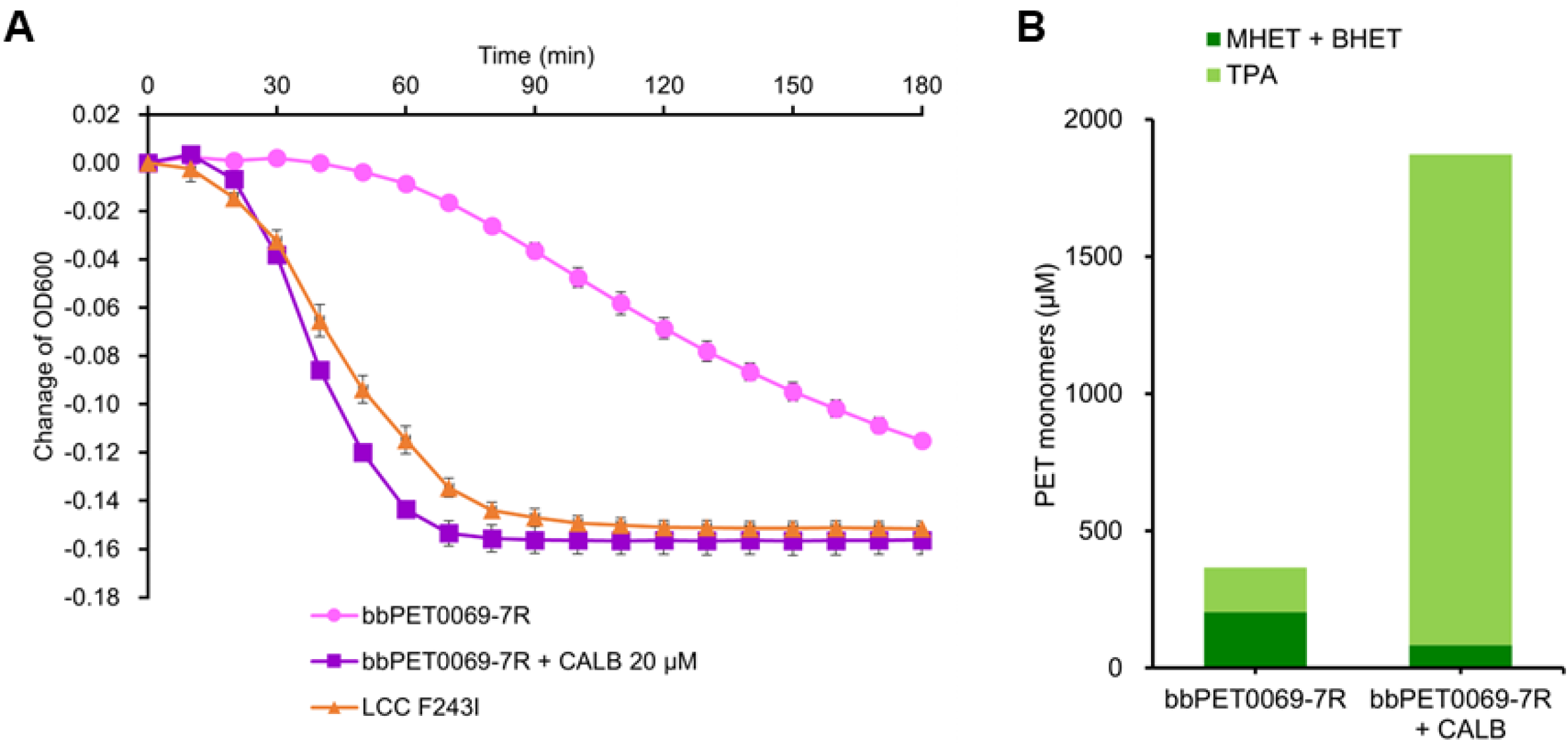
The engineered bbPET0069-7R maintained the beneficial synergy with CALB. A) bbPET0069-7R (pink), bbPET0069-7R plus CALB (purple), and LCC F243I (orange) were used for PET nanoparticle degradation assay. B) bbPET0069-7R and bbPET0069-7R plus CALB were incubated with amorphous PET film, and the released PET monomers were analyzed by HPLC.

The difference between the IsPETase engineered through directed evolution, and our approach to mutations lies in the divergent primary objectives. The previously reported mutations have primarily aimed at enhancing thermal stability, specifically raising the Tm value to get close to the Tg of PET. In contrast, our surface feature analysis concentrated mutations within the ligand-binding sites, aiming to enhance enzyme activity by altering the direct interaction between the ligand and the enzyme. In fact, among the 11 mutated sites in bbPET0069-3R, only three residues correspond to the mutated sites in ThermoPETase, DuraPETase, and HotPETase, which were Y94W, Q96Y, and R98H in bbPET0069-3R. These residues suggest a potential contribution to enhanced activity among the IsPETase mutants.

### Integration of Two-enzyme System and Mutant Enzyme

We examined whether the enhancement of PET degradation activity by CALB, (Fig. 2) could still be observed in the mutant enzymes. As a result, when CALB was added to the bbPET0069 mutants, PET degradation activity increased in all mutants (Fig. 3B, 3E). Therefore, the bbPET0069 mutants created through our directed evolution campaign retained the synergistic effect with CALB. When culturing PET nanoparticles with 0.5 µM bbPET0069-7R along with 20 µM CALB, a substantial enhancement in activity was observed, demonstrating an activity level equivalent to that of LCC F243I under the same reaction conditions (Fig. 4A). Hence, the simultaneous utilization of directed evolution and CALB effect resulted in a 12.9-fold increase in PET degradation with bbPET0069-7R compared to 12.6-fold with bbPET0069-3R from the wild-type in three hours reaction (Fig. 3B, 3E).

Furthermore, to investigate the degradation of amorphous PET film instead of PET nanoparticles, PET film degradation experiments were conducted under bbPET0069-7R + CALB conditions (Fig. 4B). As a result, the addition of CALB led to a 5.1-fold increase in the amount of Total PET monomers released from PET film, a significant decrease in the MHET and BHET, and a substantial improvement in the TPA ratio from 44.2% to 95.5%.

Based on these results, we consider the reason for the significant enhancement in PET degradation by bbPET0069 upon CALB addition as follows: CALB possesses potent BHETase and MHETase activities ^[36]^, facilitating the rapid breakdown of BHET and MHET into TPA catalyzed by CALB. Consequently, we presume that the enzymatic reaction for PET polymer and PET cleavage products significantly shifted toward the product side due to the efficient breakdown of BHET and MHET catalyzed by CALB. Recently, reports have demonstrated that adding ΔBHETase, possessing BHETase activity, improved the activity of several PET-degrading enzymes ^[19]^. BHETase and MHETase activities in PET degrading Identification of bbPET0069 as a functional PET hydrolase. A) Sequence alignment of bbPET0069 and the previously known PET hydrolases, including PHL7, LCC, IsPETase, and RgPETase. The purple box shows the G-x1-S-x2-G motif found in all PET hydrolases. Red boxes are the disulfide bonds shown in Type I and Type II. Blue boxes are the disulfide bond shown in only Type II enzymes. B) Structural alignment of the bbPET0069 structure predicted by AlphaFold2 and PHL7. C) PET hydrolase activity of bbPET0069 determined by the increase of A240. NC shows no enzyme control. D) PET degradation products (MHET, BHET, and TPA) produced by bbPET0069. NC was no enzyme control, and the level of the peaks was under the detection limit of our standard curve (labeled as the asterisk). enzyme reactions might be key functions for efficient PET degradation.

## Conclusion

In this study, we identified bbPET0069 as a new PET-degrading enzyme. The predicted enzyme structure shows similarity to cutinase-like Type I PET hydrolases but represents a new category of PET-degrading enzymes lacking disulfide bonds. Furthermore, bbPET0069 demonstrated synergistic enzymatic activity with CALB, significantly increasing PET degradation rate and total yield. Through a directed evolution, we engineered bbPET0069, and the mutant bbPET0069-3R, generated after three rounds of combining mutations identified through surface feature analysis, achieved 12.6-fold improvement in PET degradation activity compared to the wild-type enzyme. Applying the two-enzyme system utilized in this study and employing directed evolution with surface feature analysis offer a pathway to enhance newly discovered PET-degrading enzyme activity rapidly, likely to improve additional functions such as thermostability or degradation activity for crystallized PET materials in future studies.

## Acknowledgments

The acquisition of gene information and DNA synthesis were partly supported by JST FOREST JPMJFR210F.

