## Supplementation file for "A combination of two-enzyme system and enzyme engineering improved the activity of a new PET hydrolase identified from soil bacterial genome"

**Supporting Information**
**Experimental Procedures**

*Chemicals and reagents*

CALB was purchased from Novozymes as Lipozyme® CALB L. TPA, MHET, pNPB, PET film (GF51809799-1EA), hexafluoroisopropanol, and acetonitrile were purchased from Sigma-Aldrich.

*Bioinformatic analysis to identify a PET hydrolase candidate*

bit-GEM, a bacterial genome database as of 2022/4/25, was used to search DNA sequences similar to leaf compost cutinase (LCC) using MMseqs2 [37].

*Molecular cloning and E. coli transformation*

A PET hydrolase gene fragment was synthesized (Twist Biosciences) and cloned into a vector backbone DNA fragment amplified from pET-44a(+) (Novagen) by NEBuilder HiFi DNA Assembly Master Mix (New England Biolabs). The reaction mixture transformed BL21(DE3)pLysS (Merck). The DNA sequences of the PET hydrolase candidate, ORF of the transformants were verified by Sanger sequencing. The sequences of bbPET0069 and the vector backbone are listed in Table Sa.

*Protein expression and purification*

*E. coli* transformant was incubated in 3 mL of LB medium supplemented with ampicillin and chloramphenicol overnight at 37 °C. 1 mL of the preculture was transferred to 100 mL of Overnight Express™ Autoinduction System (Merck) supplemented with ampicillin and chloramphenicol and incubated overnight at 18 °C. The cells were harvested and lysed with xTractor Buffer (TaKaRa) after adding protease inhibitors. DNase I (Nippon Gene) was added to the cell lysate and incubated for 15 min at room temperature. The cell lysate was clarified by centrifugation and applied to HisTALON™ Gravity Column (TaKaRa). The HisTALON resin was washed twice with a Washing Buffer, including ten mM imidazole. C-terminally, His-tagged protein was eluted with an Elution Buffer, including 500 mM imidazole. The elution fraction was buffer exchanged to 50 mM sodium phosphate, pH 7.4, by passing the PD10 column (Cytiva). The protein concentration was assessed by BCA assay (TaKaRa). LCC and FAST-PETase were prepared as previously described [9,12].

*Esterase activity assay*

*E. coli* transformant was inoculated overnight in 3 mL of Overnight Express™ Autoinduction System supplemented with ampicillin and chloramphenicol. The cells were lysed with Lysis Buffer containing 100 mM sodium phosphate, pH 7.4, 1 g/L of lysozyme, and 0.5 g/L of colistin sulfate. 5-20 µL of the cell lysate was added to the substrate buffer containing 1X PBS, 5% acetonitrile, and one mM 4-nitrophenyl butyrate (Sigma-Aldrich). After 30 min incubation at 37 °C, the absorbance at 405 nm was measured by INFINITE M NANO+ plate reader (TECAN).

*PET film degradation assay*

Amorphous PET film (ES301445, Goodfellow) was cut into a 9 mm diameter circle with a punch. The PET film was incubated with a purified enzyme in 50 mM sodium phosphate, pH 7.4, at an appropriate concentration with vigorous shaking at 40 °C. After heat denaturation of the enzymatic activity, the total amount of PET monomers released from the PET was assessed by absorbance at 240 nm. Alternatively, the reaction supernatant was analyzed by reversed-phase HPLC (LC-2000Plus series, JASCO) using a C18 column, 4.6 x 250 mm, (Shodex) with isocratic condition using 25 % of acetonitrile/acetic acid. Peaks of terephthalic acid, MHET, and BHET were monitored with 240 nm (Supporting Fig. 5).

*PET turbidity assay*
PET nanoparticles were prepared as described previously [23]. The amorphous PET films were dissolved in hexafluoroisopropanol (Sigma-Aldrich) and precipitated in water. *E. coli* cell lysate or purified protein was mixed with the PET nanoparticle solution in a 96-well plate. The optical density of the wells was monitored by the INFINITE M NANO+ plate reader. Alternatively, absorbance at 240 nm of the supernatant was used to assess the amount of PET monomers released from PET nanoparticles. To investigate the impact of CALB on bbPET0069, we added different concentrations of CALB to the reactions. The CALB solution was buffer exchanged to 100 mM sodium phosphate pH 7.4 with a PD10 column before being used for the PET turbidity assay to reduce the background absorbance at 240 nm due to the buffer ingredients.

*Surface Feature Analysis to identify mutation site candidates of bbPET0069*
A predicted structural data of bbPET0069 was created by AlphaFold2 [21]. The predicted protein structure was visualized with PyMOL (The PyMOL Molecular Graphics System, Version 2.0 Schrödinger, LLC). Three protein surface characteristics around the binding pocket of bbPET0069 were analyzed and compared with the structural data of the existing PET hydrolases. The Surface Feature Analysis tool provided the preferable mutation sites and recommended amino acids of bbPET0069. The list of recommended mutations was used for the first and second rounds of directed evolution.

*Protein language model analysis to identify mutation site candidates of bbPET0069*
To identify preferable mutation sites of bbPET0069-3R for the fourth round of directed evolution, we performed zero-shot prediction using ESM-1v [26]. A mutation score was calculated at all amino acid residues using wt-marginals and masked-marginals. Based on the score ranking, the mutation sites of bbPET0069-3R were determined. For the sixth round of mutagenesis, ESM2 [27] was used to identify the mutated amino acids of bbPET0069-4R.

*PCR mutagenesis*
To introduce multiple mutations into the wild type or the top mutant clones obtained from the previous round of directed evolution, primers including more than one mutation site using degenerative nucleotides were designed.After 1st PCRs, the amplified DNAs were treated with DpnI and KDL mix (New England Biolabs) and used to transform DH5α cells. The plasmid DNAs were extracted from the transformed DH5α cells, and the plasmid DNA mixture was used as a template for 2nd PCRs. The 2nd PCR amplimers were treated with DpnI and KDL mix and used to transform DH5α cells. The plasmid DNAs extracted from the transformed DH5α cells were used to transform BL21(DE3)pLysS. The transformed cells were used for PET turbidity assay as described above. For the third round of the directed evolution, domain shuffling technology was used to obtain variants with different combinations of mutations. 5’-part or 3’-part of the favorable mutants obtained from the first and second rounds of directed evolution were separately amplified and randomly cloned into the vector backbone with HiFi DNA Assembly Master Mix. The resulting DNAs were used to transform BL21(DE3)pLysS. The transformed cells were used for PET turbidity assay to assess their PET hydrolase activity.

**Supporting figures**

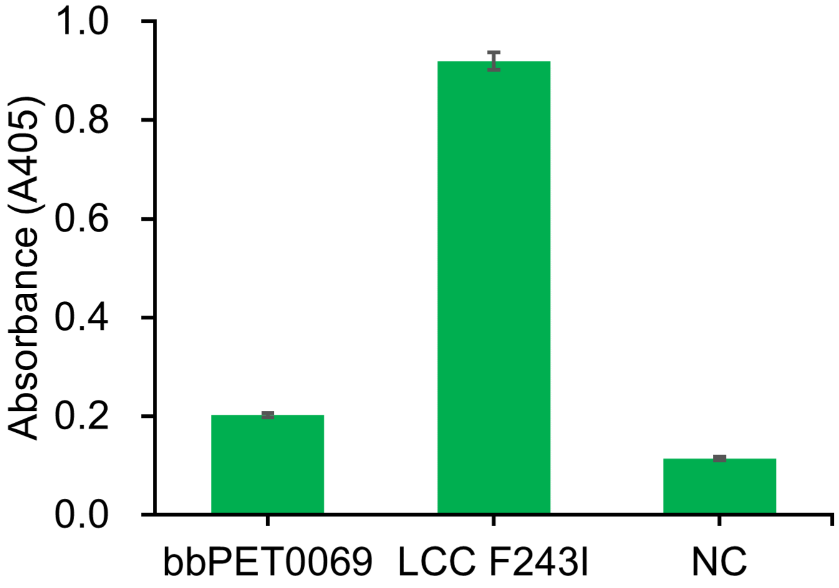


**Supporting Fig. 1.** Esterase activity assay with pNPB substrate. The cell lysate, which expresses bbPET0069 or LCC F243I, and NC (empty vector control) were used for this assay.

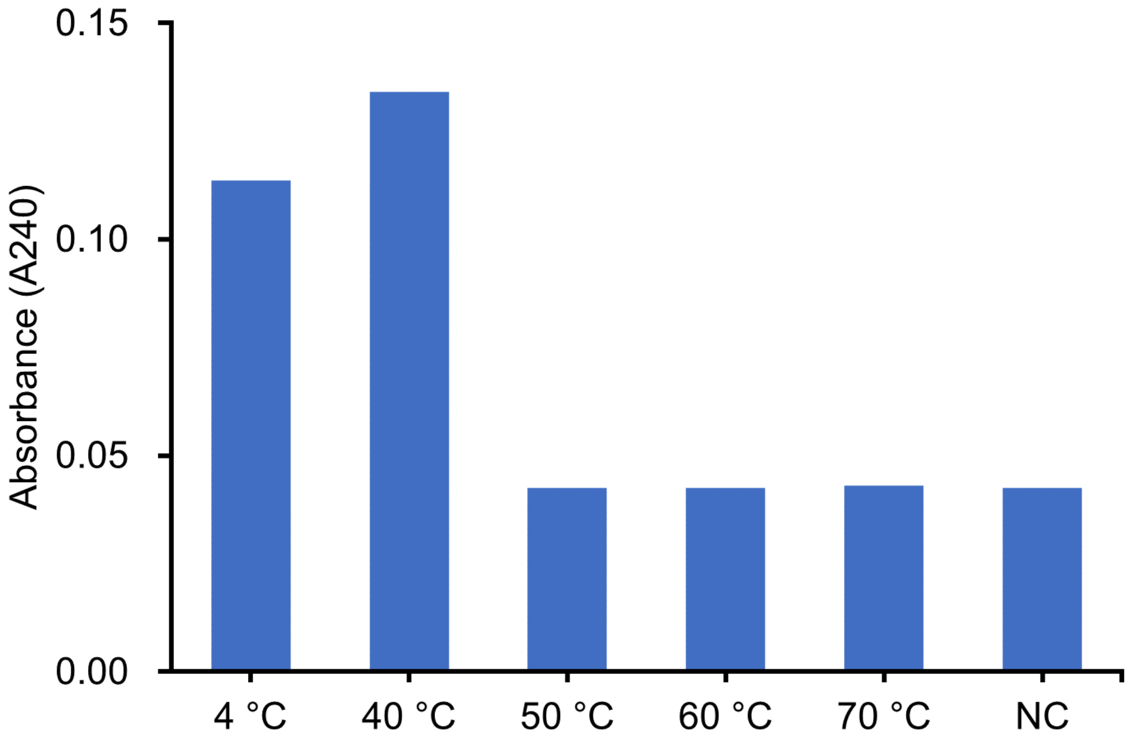


**Supporting Fig. 2.** Esterase activity of bbPET0069 preincubated with each temperature. NC is an empty vector control.

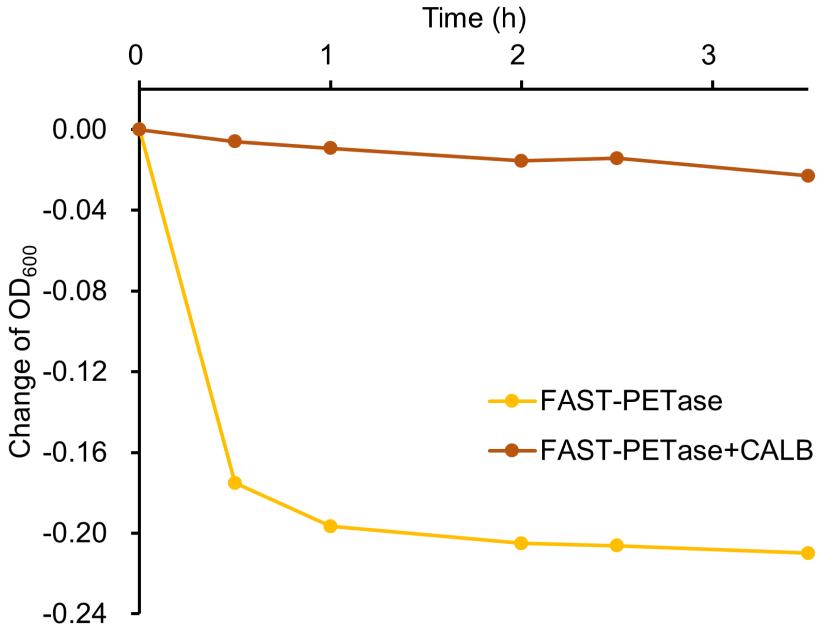


**Supporting Fig. 3.** PET nanoparticle degradation assay with FAST-PETase (yellow) and FAST-PETase plus CALB (brown).

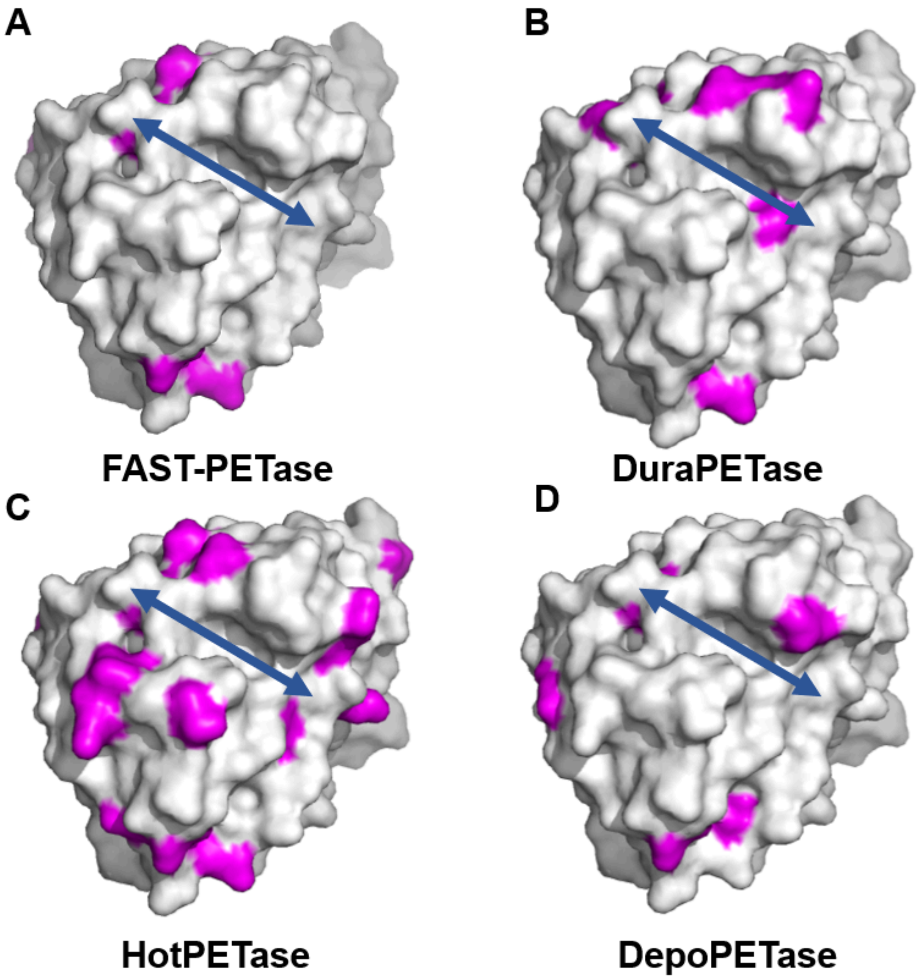

**Supporting Fig. 4.** The mutated amino acids in four different engineered PETases. FAST-PETase (A), DuraPETase (B), HotPETase (C), and DepoPETase (D). The mutated amino acids were labeled with purple on the crystal structure of IsPETase.

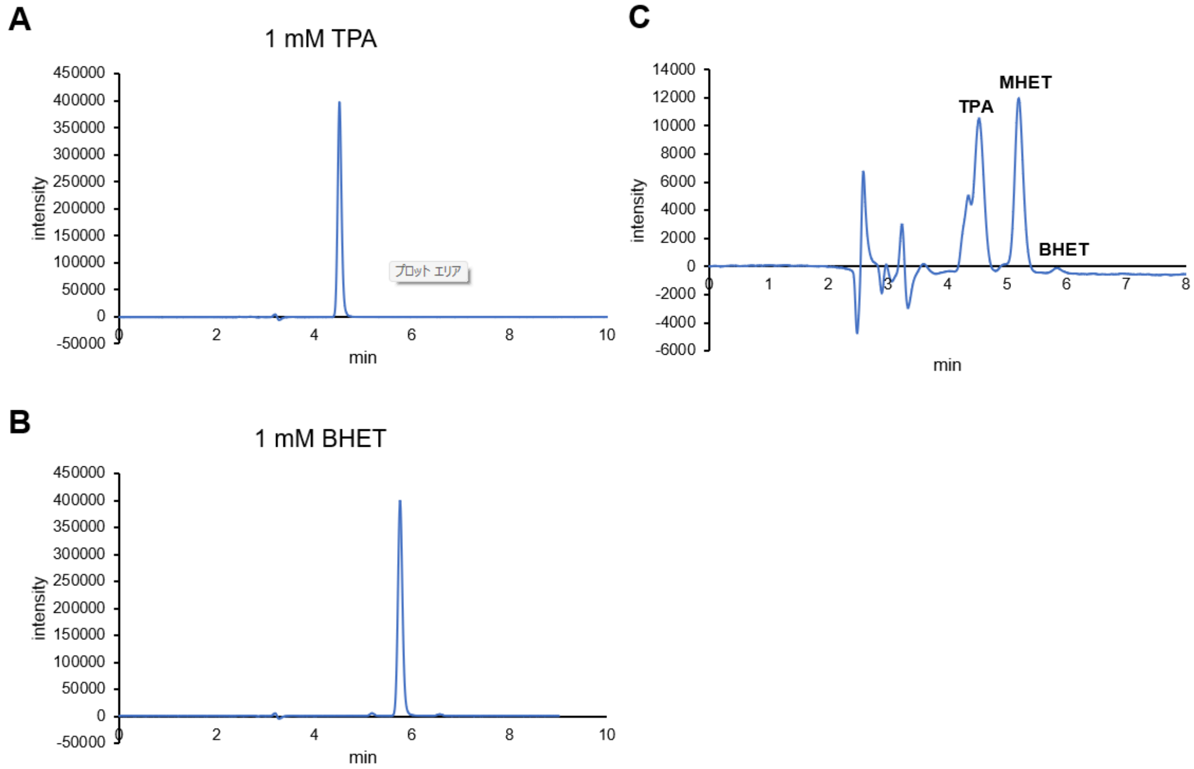


**Supporting Fig. 5.** HPLC profile of TPA (A), BHET (B), and the reaction solution incubated with PET film and bbPET0069 (C). HPLC was performed as described in the Materials and Methods section.

**Supporting Table 1**

Supporting Table 1. A mutagenesis plan used for each round of the directed evolution campaign.

| No. of muation round | A method to identify the mutation sites |
|---|---|
| 1st round | Surface Feature Analysis |
| 2nd round | Surface Feature Analysis |
| 3rd round | Shuffling of the mutants obtained through 1st and 2nd rounds |
| 4th round | Protein Language Model (ESM-1v) |
| 5th round | Combination of 4th round mutations |
| 6th round | Protein Language Model (ESM2) |
| 7th round | Active site mutations of known PET hydrolases |

**Supporting Table 2**

Supporting Table 2. Mutation sites and total mutation numbers found in all mutation rounds.

| Enzymes | Mutation | Mutation number (vs WT) |
|---|---|---|
| bbPET0069-1R | S76D | 1 |
| bbPET0069-2R | S76P/R93K/Y94W/Q96T/R98F | 5 |
| bbPET0069-3R | Y64F/S76P/R93K/Y94W/Q96Y/R98H/T179K/V180L/S182P/S209N/F211L | 11 |
| bbPET0069-4R | Y64F/S76P/R93K/Y94W/Q96Y/R98H/M112S/T179K/V180L/S182P/T195P/Q196H/E198P/Q199A/S209N/F211L | 16 |
| bbPET0069-5R | R40G/Y64F/S76P/R93K/Y94W/Q96Y/R98H/M112S/V116A/V117L/P119N/W163F/T179K/V180L/S182P/T195P/Q196H/E198P/Q199A/S209N/F211L/A247D/V248T/G249S/Y250S | 25 |
| bbPET0069-6R | I26V/Q28E/K54T/Y64F/S76P/R93K/Y94W/Q96Y/R98H/M112S/T179K/V180L/S182P/T195P/Q196H/E198P/Q199A/S209N/F211L | 19 |
| bbPET0069-7R | I26V/Q28E/K54T/Y64F/S76P/R93K/Y94W/Q96Y/R98H/M112S/T179K/V180L/S182P/R184S/T195P/Q196H/E198P/Q199A/S209N/F211L | 20 |

### Supporting Table 3

Supporting Table 3. Amino acids found in each catalytic triad, subsite I, subsite II, and extended loop region。

| Type | Enzyme | Catalytic triad | | | Subsite I | | | | | | | Subsite II | | | | | Extended loop region | | | | | | | | | |
|---|---|---|---|---|---|---|---|---|---|---|---|---|---|---|---|---|---|---|---|---|---|---|---|---|---|---|
| I | bbPET0069 | S132 | D178 | H210 | F64 | M133 | W157 | V180 | Y94 | Q96 | H186 | F190 | T65 | A66 | S70 | H131 | F211 | A212 | P213 | N214 | V215 | S216 | N217 | - | - | - | A175 |
| I | bbPET0069-3R | S132 | D178 | H210 | Y64 | M133 | W157 | L180 | W94 | Y96 | H186 | F190 | T65 | A66 | S70 | H131 | L211 | A212 | P213 | N214 | V215 | S216 | N217 | - | - | - | A175 |
| I | PHL7 | S131 | D177 | H209 | F63 | M132 | W156 | I179 | L93 | Q95 | H185 | F189 | T64 | A65 | S69 | H130 | L210 | V211 | S212 | N213 | T214 | P215 | D216 | - | - | - | A174 |
| I | LCC | S165 | D210 | H242 | Y95 | M166 | W190 | V212 | F125 | Y127 | H218 | F222 | T96 | A97 | S101 | H164 | F243 | A244 | P245 | N246 | S247 | N248 | N249 | - | - | - | A207 |
| IIb | IsPETase | S160 | D206 | H237 | Y87 | M161 | W185 | I208 | L117 | Q119 | S214 | I218 | T88 | A89 | S93 | W159 | S238 | C239 | A240 | N241 | S242 | G243 | N244 | S245 | N246 | Q247 | C203 |

### Sequences

Amino acid sequences of the PET degrading enzymes.

>bbPET0069
MADNPFERGPNPSESLIEQRRGNYDIAQRSISRFSSDGFRDGTIYHPTDTSDGKFGVIAI
SPGYTAGESSIAWLGSRIASFGFVVVTINTTTRYDQPRQRSTQLLAALDHAMNDSVVGPL
IDPERQAVMGHSMGGGGALQAAESRPEIDAVVALTPWNLKKNWDGVDAATLIIGAERDTV
ASVRTHSIPFYESLTQAEQRGYLELRGASHFAPNVSNTTIAKYSIAWMKRYLDNDDRYDQ
FLNPGPAVGYASGVSDYRLQHHHHHH

>bbPET0069-1R
MADNPFERGPNPSESLIEQRRGNYDIAQRSISRFSSDGFRDGTIYHPTDTSDGKFGVIAI
SPGYTAGESSIAWLGDRIASFGFVVVTINTTTRYDQPRQRSTQLLAALDHAMNDSVVGPL
IDPERQAVMGHSMGGGGALQAAESRPEIDAVVALTPWNLKKNWDGVDAATLIIGAERDTV
ASVRTHSIPFYESLTQAEQRGYLELRGASHFAPNVSNTTIAKYSIAWMKRYLDNDDRYDQ
FLNPGPAVGYASGVSDYRLQHHHHHH

>bbPET0069-2R
MADNPFERGPNPSESLIEQRRGNYDIAQRSISRFSSDGFRDGTIYHPTDTSDGKFGVIAI
SPGYTAGESSIAWLGPRIASFGFVVVTINTTTKWDTPFQRSTQLLAALDHAMNDSVVGPL
IDPERQAVMGHSMGGGGALQAAESRPEIDAVVALTPWNLKKNWDGVDAATLIIGAERDTV
ASVRTHSIPFYESLTQAEQRGYLELRGASHFAPNVSNTTIAKYSIAWMKRYLDNDDRYDQ
FLNPGPAVGYASGVSDYRLQHHHHHH

>bbPET0069-3R
MADNPFERGPNPSESLIEQRRGNYDIAQRSISRFSSDGFRDGTIYHPTDTSDGKFGVIAI
SPGFTAGESSIAWLGPRIASFGFVVVTINTTTKWDYPHQRSTQLLAALDHAMNDSVVGPL
IDPERQAVMGHSMGGGGALQAAESRPEIDAVVALTPWNLKKNWDGVDAATLIIGAERDKL
APVRTHSIPFYESLTQAEQRGYLELRGANHLAPNVSNTTIAKYSIAWMKRYLDNDDRYDQ
FLNPGPAVGYASGVSDYRLQHHHHHH

>bbPET0069-4R
MADNPFERGPNPSESLIEQRRGNYDIAQRSISRFSSDGFRDGTIYHPTDTSDGKFGVIAI
SPGFTAGESSIAWLGPRIASFGFVVVTINTTTKWDYPHQRSTQLLAALDHASNDSVVGPL
IDPERQAVMGHSMGGGGALQAAESRPEIDAVVALTPWNLKKNWDGVDAATLIIGAERDKL

APVRTHSIPFYESLPHAPARGYLELRGANHLAPNVSNTTIAKYSIAWMKRYLDNDDRYDQ
FLNPGPAVGYASGVSDYRLQHHHHHH

>bbPET0069-5R
MADNPFERGPNPSESLIEQRRGNYDIAQRSISRFSSDGFGDGTIYHPTDTSDGKFGVIAI
SPGFTAGESSIAWLGPRIASFGFVVVTINTTTKWDYPHQRSTQLLAALDHASNDSALGNL
IDPERQAVMGHSMGGGGALQAAESRPEIDAVVALTPWNLKKNFDGVDAATLIIGAERDKL
APVRTHSIPFYESLPHAPARGYLELRGANHLAPNVSNTTIAKYSIAWMKRYLDNDDRYDQ
FLNPGPDTSSASGVSDYRLQHHHHHH

>bbPET0069-6R
MADNPFERGPNPSESLIEQRRGNYDVAERSISRFSSDGFRDGTIYHPTDTSDGTFGVIAI
SPGFTAGESSIAWLGPRIASFGFVVVTINTTTKWDYPHQRSTQLLAALDHASNDSVVGPL
IDPERQAVMGHSMGGGGALQAAESRPEIDAVVALTPWNLKKNWDGVDAATLIIGAERDKL
APVRTHSIPFYESLPHAPARGYLELRGANHLAPNVSNTTIAKYSIAWMKRYLDNDDRYDQ
FLNPGPAVGYASGVSDYRLQHHHHHH

>bbPET0069-7R
MADNPFERGPNPSESLIEQRRGNYDVAERSISRFSSDGFRDGTIYHPTDTSDGTFGVIAI
SPGFTAGESSIAWLGPRIASFGFVVVTINTTTKWDYPHQRSTQLLAALDHASNDSVVGPL
IDPERQAVMGHSMGGGGALQAAESRPEIDAVVALTPWNLKKNWDGVDAATLIIGAERDKL
APVSTHSIPFYESLPHAPARGYLELRGANHLAPNVSNTTIAKYSIAWMKRYLDNDDRYDQ
FLNPGPAVGYASGVSDYRLQHHHHHH

>LCC F243I
MKYLLPTAAAGLLLLAAQPAMAQSNPYQRGPNPTRSALTADGPFSVATYTVSRLSVSGFG
GGVIYYPTGTSLTFGGIAMSPGYTADASSLAWLGRRLASHGFVVLVINTNSRFDYPDSRA
SQLSAALNYLRTSSPSAVRARLDANRLAVAGHSMGGGGTLRIAEQNPSLKAAVPLTPWHT
DKTFNTSVPVLIVGAEADTVAPVSQHAIPFYQNLPSTTPKVYVELDNASHIAPNSNNAAI
SVYTISWMKLWVDNDTRYRQFLCNVNDPALSDFRTNNRHCQHHHHHH

>FAST-PETase
MAQTNPYARGPNPTAASLEASAGPFTVRSFTVSRPSGYGAGTVYYPTNAGGTVGAIAIVP
GYTARQSSIKWWGPRLASHGFVVITIDTNSTLDQPESRSSQQMAALRQVASLNGTSSSPI
YGKVDTARMGVMGWSMGGGGSLISAANNPSLKAAAPQAPWHSSTNFSSVTVPTLIFACEN
DSIAPVNSSALPIYDSMSQNAKQFLEIKGGSHSCANSGNSNQALIGKKGVAWMKRFMDND
TRYSTFACENPNSTAVSDFRTANCSHHHHHH
